# Late mitochondrial origin is pure artefact

**DOI:** 10.1101/055368

**Authors:** William F. Martin, Mayo Roettger, Chuan Ku, Sriram G. Garg, Shijulal Nelson-Sathi, Giddy Landan

## Abstract

Pittis and Gabaldón^1^ recently claimed that the mitochondrion came late in eukaryotic evolution, following an earlier phase of evolution in which the eukaryotic host lineage acquired genes from bacteria. Here we show that their paper has multiple fatal flaws founded in inappropriate statistical methods and analyses, in addition to erroneous interpretations.

---

For 1,078 phylogenetic trees containing prokaryotic and eukaryotic homologues, Pittis and Gabaldon^1^ calculated the length of the branch subtending the eukaryotic clade (raw stem length, *rsl*) relative to the median root-to-tip length of lineages within the eukaryotic clade (eukaryotic branch length, *ebl_med_*), a value they call stem length (*sl*). From variation in *sl*, they infer early (large *sl*) and late (small *sl*) gene acquisitions in eukaryotes, using *sl* as a measure for age. They feed values of *sl* into the expectation maximization (EM) algorithm to obtain a fit composed of five Gaussians, one component containing 14 very large values, which they exclude from further analysis. The remaining 1,064 values of *sl* are sorted into four components, analyses of which they interpret as evidence that some genes entered the eukaryote lineage early (component 4), some later (component 3), some later yet (component 2) and the largest portion finally entering with the mitochondrion (component 1).

The first question is: Are these four components real? No. They are an artefact produced by the over-fitting of a complex (14 parameters) Gaussian mixture model, when a much simpler (2 parameters) log-normal model better explains the data. The *sl* data of Pittis and Gabaldón, which we show in Fig.1a for inspection, are not multiple Gaussian distributed with five components, they are log-normally distributed, as borne out by both the Akaike and the Bayesian information criteria (Fig. 1b). This is the cardinal fatal flaw of Pittis and Gabaldón^1^. Their four (five-exclude-one) Gaussian groups are a methodological artefact. All analyses, tests and far-reaching inferences about eukaryote origin based upon the four Gaussian mixture components^1^ of *sl* are not just erroneous, they are meaningless, because the data are not normally distributed, with five components or otherwise.

How do they obtain a five-component mixture model for *sl*? They incorrectly treat the *sl* values as normally distributed. The *sl* values are ratios, hence *strictly positive*, with mean 0.48, standard deviation (SD) 0.54, and skewness 4.7. Because negative values are within one SD from the mean, and because the distribution is not symmetrical, the *sl* values cannot possibly be normally distributed. For data with such features, a logarithmic transformation is to be examined^2^. The transformed *sl* values do fit a Gaussian, that is, the *sl* values should be modeled by a log-normal distribution. Elementary statistical procedures were neglected, and since one Gaussian did not fit the data, more Gaussians were needlessly presumed^1^. This is a textbook case of over-fitting, where the addition of new parameters increases the apparent fit (Fig.1b), even when the underlying model is inappropriate. The EM programme reproducibly generates 3–5 Gaussian components from randomly generated, perfectly log-normal data (see Methods) of the sample size, mean and variance reported^1^.

Their partitioning of the data into four components, the central pillar of their paper, is thus fatally flawed. But so is the use of *sl* values to draw inferences about evolutionary time. Since different gene families evolve at different rates, the raw *rsl* distances are normalized by *ebl_med_*, which is claimed to reflect, for each gene family, a characteristic eukaryotic evolutionary rate that was constant across all lineages and times during eukaryotic evolution: a root-to-tip molecular clock for each tree. A clock assumption might hold for some gene families^3^, but it does not hold for the majority of the 1,078 families reported^1^. The full set of *ebl* values for each gene family reveals extreme variation, with a mean per-family coefficient of variation of 27%, and a median longest-to-shortest within family *ebl* ratio of 2.2. Across their 1,078 trees^1^, the largest value of *ebl* exceeds that of the shortest by >2-fold — on average. Clearly, the molecular clock assumption is not met, and *ebl_med_* is neither characteristic nor constant (Fig. 1c).

**Figure. 1.**
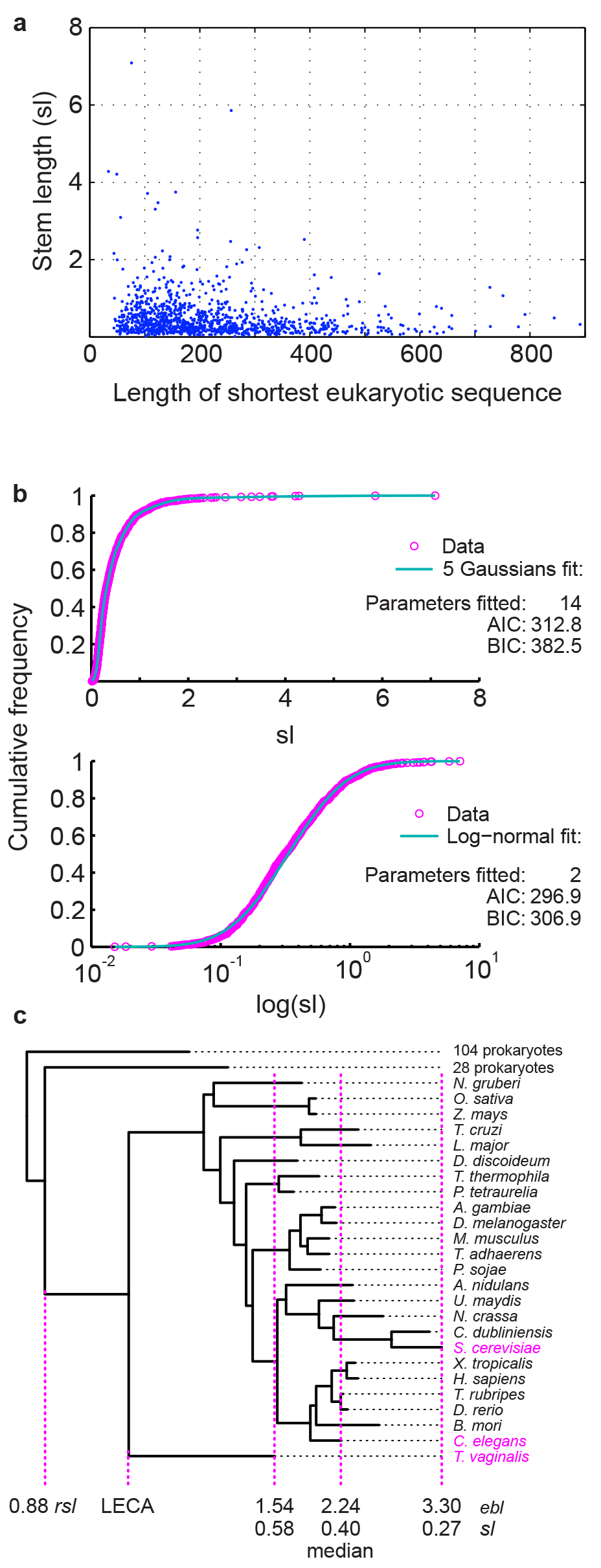
Distribution of 1,078 stem length (*sl*) values. (a) *sl* as a function of sample size (eukaryotic sequence length)
(b) Fit of the *sl* values to a five Gaussian mixture model (top), and to a log-normal model (bottom). AIC: Akaike information criterion, BIC: Bayesian information criterion. Note that the log-normal distribution is strongly preferred.
(c) Phylogenetic tree and *sl* derivation for COG4178_01, an ABC transporter present in 25 eukaryotic taxa. Which eukaryotic branch length (*ebl*) should be used to calibrate the raw stem length (*rsl*)? The minimal, median and maximal lineages are highlighted in magenta. Perchance it is a moot question, as in the absence of a LECA-to-present molecular clock, none of the resulting *sl* values convey meaningful information. The ratio of longest to shortest *ebl* is 2.2 (2.15 unrounded), a value representative of the dataset as 579 other trees have larger ratios.

Dividing *rsl* by *ebl_med_* to produce *sl* is then bound to yield arbitrary values, which it does, and the interpretation of these values as measures of divergence times culminates in absurd results. How so?

Eukaryotes are at least 1.6-1.8 billion years (Ga) in age^4^. If one uses *sl* as a measure for the age of genes that eukaryotes acquired from prokaryotes^1^, variation in *sl* implies continuous eukaryotic gene acquisition from prokaryotes starting >4.5 Ga ago^1^, before Earth's formation. That seems unlikely. Where is the error? Examining values of *sl* for groups within eukaryotic phylogeny are instructive. Crucially, all well-sampled eukaryotic groups show variation and distribution of *sl* virtually identical to that of eukaryotes as a whole (Fig. 2). The log-normal distribution again fits the data best, yet it is all-too-easy to use EM to over-fit a Gaussian mixture model with multiple components. Does this imply phases of early and late acquisition of genes from other eukaryotes? For example, the value of *sl* for metazoans, as defined^1^, indicates the age of the metazoan stem lineage after divergence from other eukaryotes relative to the age of the metazoan crown. Taking the crown age of metazoans as ~1 Ga^4^, the metazoan stem lineage, with *sl* ranging from ~0.1 to ~3, diverged continuously *from its eukaryotic sistergroup* during the time ~0.1 Ga to ~3 Ga before the first metazoan arose ~1 Ga ago, which cannot be true^4^,^5^. We have a far less radical alternative explanation: *sl* is not an indicator of gene age differences within or between trees at all, rather *sl* vividly documents abundant branch length variation within and among Pittis and Gabaldon's trees, stemming from rate variation within and among lineages across trees, which is well-known to exist, which is expected^3^^4^^6^, and which can be readily grasped by looking at actual trees (Fig. 1c).

In addition, their 1,078 trees^1^ are not independent samples of the data. Starting from 883 EggNOG clusters, 722 clusters were used once, 130 twice, 28 thrice, and 3 clusters in four trees. Trees showing eukaryote polyphyly were split and scored as multiple eukaryote monophyly^1^. Their 1,078 trees contain 403,451 sequences: 238,080 occur once, 5 occur in seven trees, 3 in six, 53 in five, 2318 in four, 14,645 in three, and 55,923 sequences occur in two different trees. Moreover, their statistical analysis of α-proteobacterial *versus* bacterial but non-α- proteobacterial gene classes hinges upon rare and/or anomalous data: if alignments containing very short, highly gapped or otherwise tenuous attributes are removed, or if analyses are properly restricted to their 722 independent samples, their borderline significance values suggesting two classes disappear completely (Fig. 3).

**Figure. 2.**
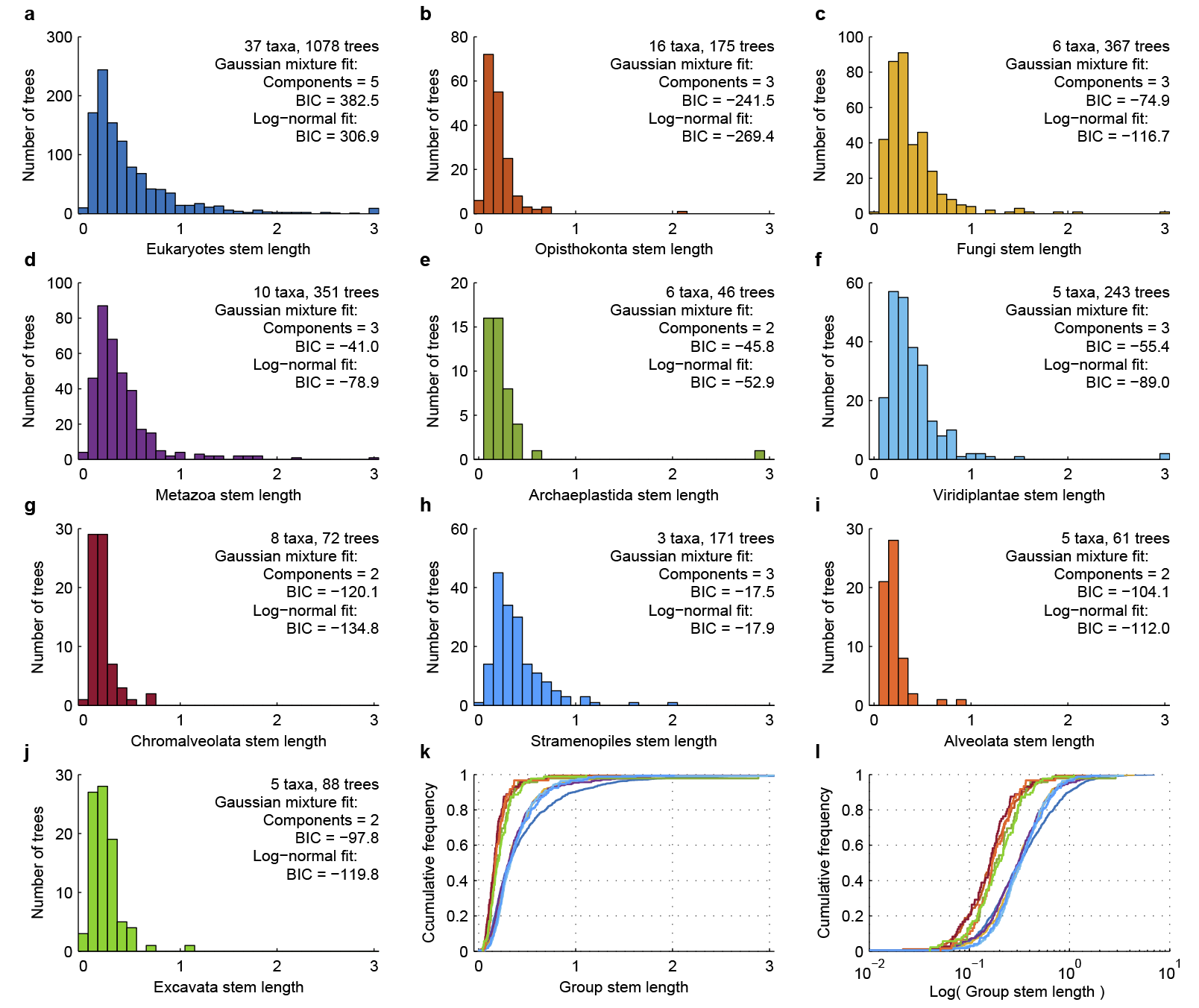
Stem length (*sl*) distributions among eukaryotes and the fit to Gaussian mixture and log-normal models. (a-j) Histograms of group specific *sl* for the largest clade containing only group members with taxa from at least two taxonomic sub-groups. Values in panel (a) are from reference 1, values in panels (b-j) were calculated from the trees in reference 1. In panels (a-j), the rightmost bin contains all values ≥3; AIC: Akaike information criterion, BIC: Bayesian information criterion. (k-l) Empirical cumulative distribution functions for the *sl* values in panels (a-j), in *sl* scale (k) and *log(sl)* scale (l). Colors match the colors used in (a-j).

Unnoted by Pittis and Gabaldón^1^, an earlier study analyzed more than three times as many independent trees^7^. In that study, all sequences were unique, eukaryote non-monophyly was scored as such^7^, not as multiple observations of eukaryote monophyly^1^, and the data uncovered neither evidence for a late mitochondrion^7^, nor for a late plastid^7^.

In summary, *sl*-based conclusions about eukaryote evolution^1^ are unfounded, resting upon fatal flaws in i) over-fitting of the wrong distribution model, ii) analyses of non-independent data, and iii) implicit, untested, and untrue molecular clock assumptions. Some journals require authors to document the appropriateness of their statistics and methods at the submission stage^8^. For the paper by Pittis and Gabaldon^1^, that apparently did not occur, possibly sending the wrong signal to young scientists and the community that the improper use of statistical methods is acceptable if one obtains a particular result.

**Figure. 3.**
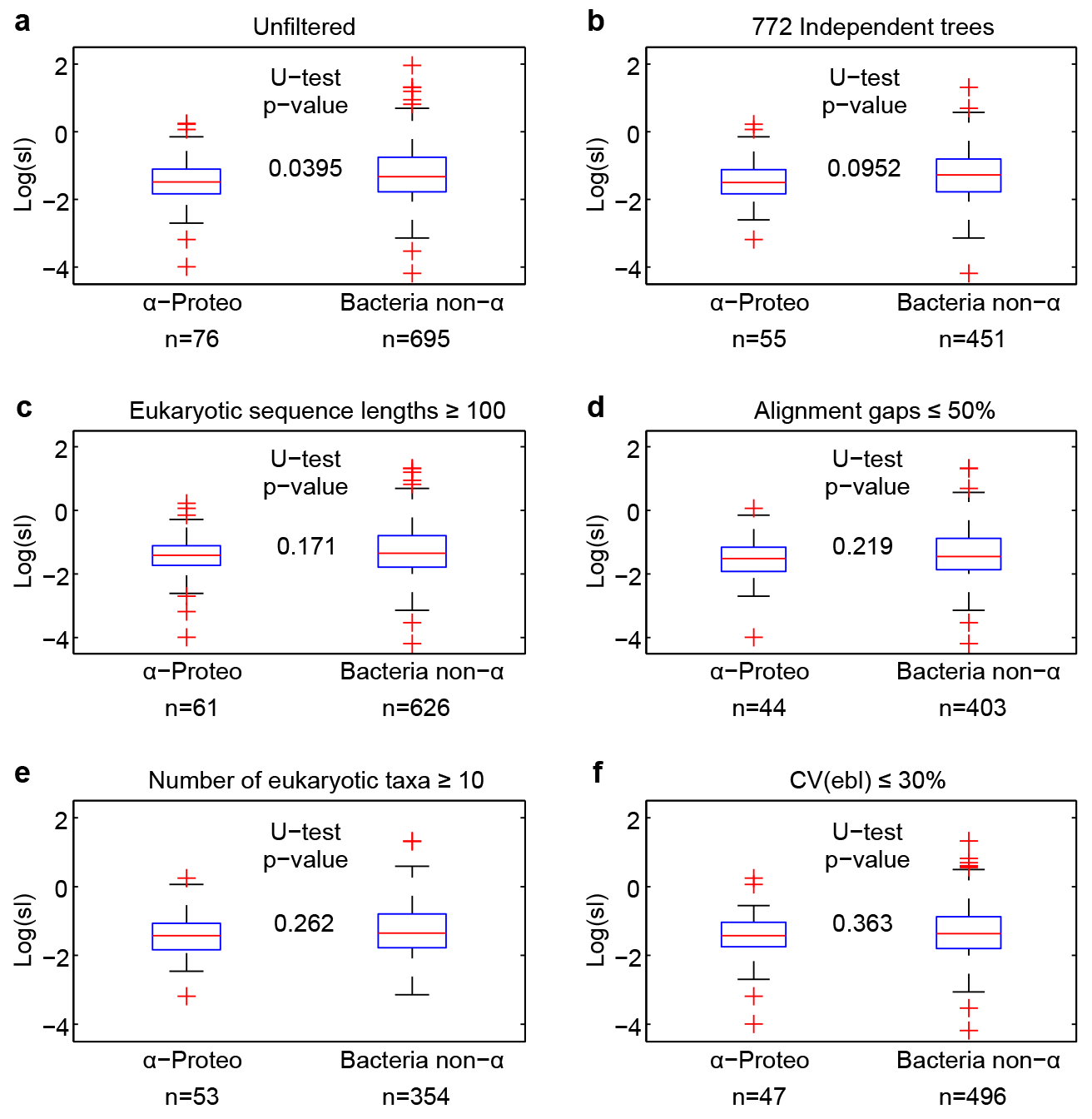
Comparison of stem length (*sl*) values in classification of the prokaryotic sister clade as α-proteobacterial or bacterial but non-α-proteobacterial. (a) Unfiltered: full dataset analyzed in reference 1. (b-f) Datasets obtained by exclusion of questionable, low-quality, or non-independent sample points. n: number of observations, U-test: Mann-Whitney U test, CV: Coefficient of variation.

## Methods

All analyses were based on alignments and phylogenetic trees kindly provided by T. Gabaldón. No re-alignments or re-inference of trees was carried out. Values of *rsl* and *ebl* were extracted from the trees, values of *sl* were recalculated, reproducing the values reported^1^. For calculating *sl* within eukaryotic groups, trees were searched for the largest clade containing only group members with taxa from at least two different taxonomic sub-groups. All statistical analyses were performed using the MatLab^®^ statistics toolbox.

## Author contributions

W.F.M., MR., C.K., S.G.G., S.N.-S. and G.L designed experiments, analyzed data and prepared this manuscript; M.R., C.K., S.N.-S. and G.L performed computational analysis.

